## Supplementary for "Δ133p53α and Δ160p53α isoforms of the tumor suppressor protein p53 exert dominant-negative effect primarily by co-aggregation"

**Supplementary Table 1. Sequences of oligonucleotides used in the work.**

| Name | Sequence 5' to 3' | Description | Source |
|---|---|---|---|
| **Primers for construction of Δ133p53 and Δ160p53 with FLAG-tag expression plasmids** | | | |
| Δ133p53-FLAG fw | CTAGCTAGCCACCATGTTTTGCCAACTGGCCAAG | Δ133p53-FLAG | This work |
| Δ160p53-FLAG fw | CTAGCTAGCCACCATGGCCATCTACAAGCAGTCA | Δ160p53-FLAG | This work |
| Δ133/160p53-FLAG rev | CCGGAATTCTTATTTATCGTCATCGTC | Δ133/160p53- FLAG | This work |
| **Primers for construction of FLp53 and its two isoforms with V5-tag expression plasmids** | | | |
| FLp53-V5 fw | CCGGAATTCGCCACCATGGAGGAGCC | FLp53-V5 | This work |
| FLp53-V5 rev | CCGCTCGAGGTCTGAGTCAGGCCCTTCTG | FLp53-V5 | This work |
| Δ133p53-V5 fw | CCGGAATTCGCCACCATGTTTTGCCAACTGGCCAAG | Δ133p53-V5 | This work |
| Δ160p53-V5 fw | CCGGAATTCGCCACCATGGCCATCTACAAGCAG | Δ160p53-V5 | This work |
| **Primers for construction of untagged proteins expression plasmids** | | | |
| FLp53 fw | CTAGCTAGCCACCATGGAGGA | FLp53 | This work |
| FLp53 rev | CCGGAATTCTTAGTCTGAGTCAGGCCCTTCT | FLp53 | This work |
| Δ133p53 fw | CTAGCTAGCCACCATGTTTTGCCAACTGGCCAAG | Δ133p53 | This work |
| Δ160p53 fw | CTAGCTAGCCACCATGGCCATCTACAAGCAG | Δ160p53 | This work |
| **Primers for construction of Bax-Luc expression plasmids** | | | |
| baxP fw | CCGCTCGAGGCTTCAGCCCGGGAAT | BAX promoter | This work |
| baxP rev | CCCAAGCTTAGCTCTCCCCAGCGCAGA | BAX promoter | This work |
| **Primers for ChIP-qPCR assay** | | | |
| p21 5`RE fw | AGCAGGCTGTGGCTCTGATT | p53 5`RE of p21 promoter | [1] |
| p21 5`RE rev | CAAAATAGCCACCAGCCTCTTCT | p53 5`RE of p21 promoter | [1] |
| MDM2 fw | TCAAGTTCAGACACGTTCCGAA | p53 RE of MDM2 promoter | [1] |
| MDM2 rev | CTGGGAAAATGCATGGTTTAAATA | p53 RE of MDM2 promoter | [1] |
| PUMA fw | TCAGTGTGTGTGTCCGACTGTC | p53 RE of PUMA promoter | [1] |
| PUMA rev | GGCAGGGCCTAGCCCA | p53 RE of PUMA promoter | [1] |
| Bax fw | AGGCTGAGACGGGGTTATCT | p53 RE of BAX promoter | This work |

| *Continued* | | | |
|---|---|---|---|
| Bax rev | CAAGTGCAAAAGCTCAGAGG | p53 RE of BAX promoter | This work |

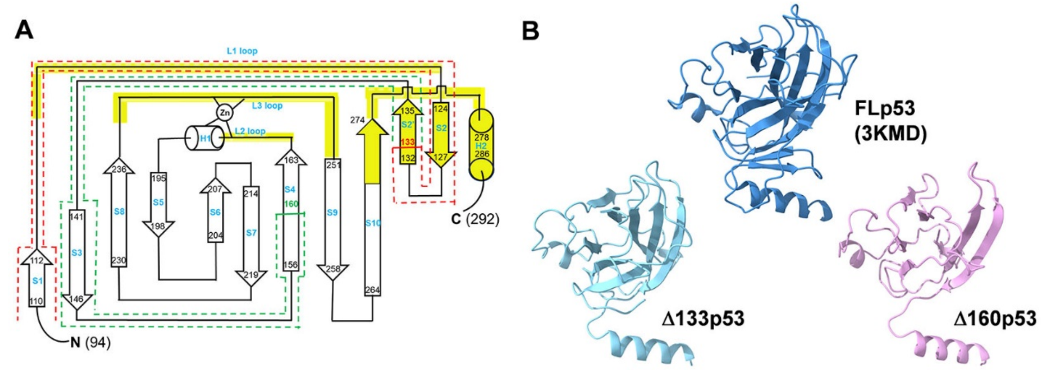


**Supplementary Figure 1. Core domain structures of full-length (FL) p53 and its isoforms Δ133p53 and Δ160p53.**

**(A)** Topology diagram of the p53 DNA-binding domain based to prior researches[2, 3]. Residues S94 to K132 are outlined with red dash rectangles. Furthermore, the residues M133 to A159 are outlined with green dash rectangles. The β-strands (S), $\alpha$ helices (H), loops (L), and the zinc atom (Zn) are labeled. The DNA-binding surface is emphasized in yellow and includes two large loops (L2 and L3) and the loop-sheet-helix (LSH) motif (L1, β-hairpin S2–S2′, H2 and the C-terminal residues of S10). **(B)** Comparison of core domain structures. The core domain structures of FLp53 (A chain, residues 92–291, PDB ID: 3KMD) and its isoforms Δ133p53 (residues 133-291, predicted by Alphafold2) and Δ160p53 (residues 160-291, predicted by Alphafold2[4]). Each structure is presented in the same orientation to facilitate direct comparison. The structures are analyzed by ChimeraX[5, 6].

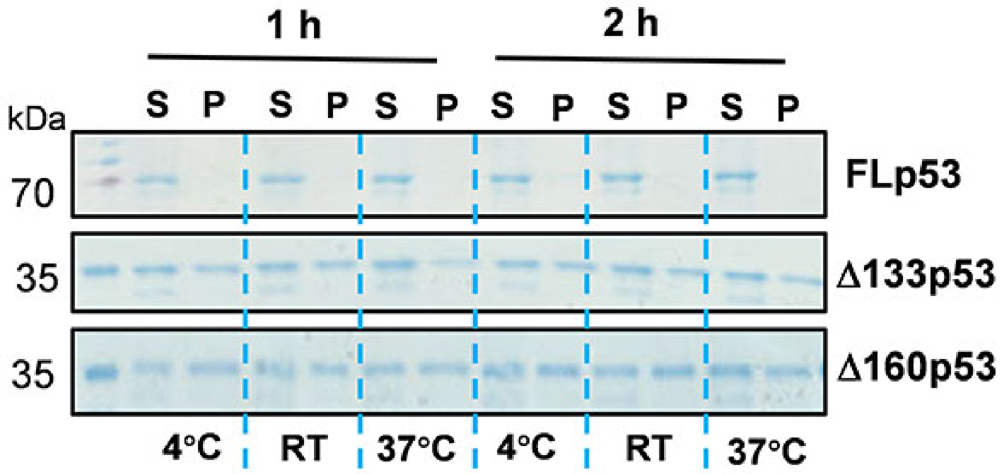


**Supplementary Figure 2. Aggregation propensity of FLp53 and its Δ133p53 and Δ160p53 isoforms by sedimentation analysis in SDS-PAGE.** Purified proteins FLp53 and its isoforms Δ133p53 and Δ160p53 at a concentration of 1 μM incubated in 20 μL of reaction buffer (20 mM Tris-HCl pH 7.9, 150 mM NaCl, 5 mM DTT, 5% glycerol) at 4°C, room temperature (RT) (approximately 23°C), or 37°C for 1 h or 2 h. After incubation, the samples were centrifuged at 20,000 × g for 30 min. The pellets were resuspended in 20 μL buffer. Both the supernatant (S) and pellet (P) fractions were analyzed by SDS-PAGE. While the FLp53 protein was detected only in the soluble supernatent fraction irrespective of temperature, a significant amount of Δ133p53 and Δ160p53 proteins were detected in the insoluble pellet fraction. This result suggests that the Δ133p53 and Δ160p53 isoforms are inherently destabilized and have much higher aggregation propensity than FLp53. FLp53 protein was sourced from Sigma (catalog number P6247), while Δ133p53 and Δ160p53 proteins were purified in our lab using multiple column chromatography.

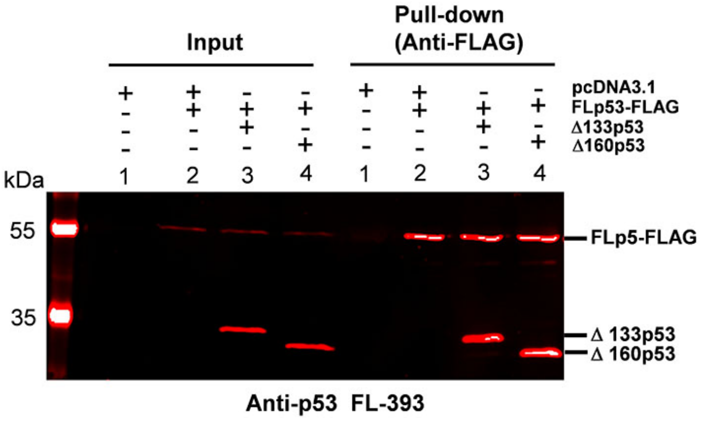


**Supplementary Figure 3. Hetero-oligomers formation between FLp53 and its isoforms Δ133p53 or Δ160p53.** Co-immunoprecipitation of FLAG tagged FLp53 with untagged Δ133p53 and Δ160p53 in H1299 cells. The complexes were isolated with anti-FLAG antibody-coupled magnetic beads. Western blot analysis was performed using the anti-p53 FL-393 antibody, which targets amino acids 1-393 corresponding to the full-length p53 protein. Input corresponds to 5% of the whole cell lysate.

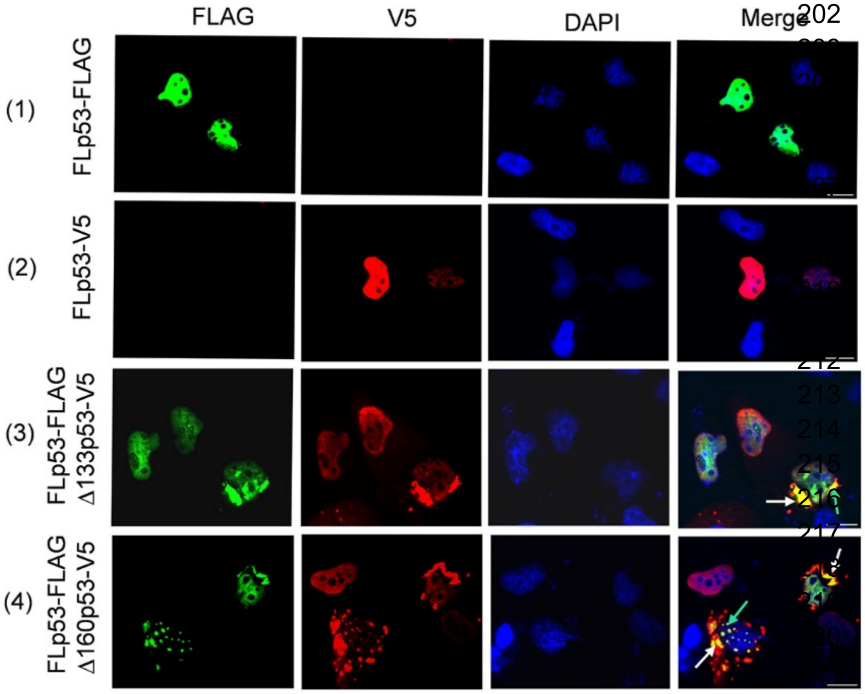


**Supplementary Figure 4. Immunofluorescence analysis of FLp53 and its isoforms Δ133p53 and Δ160p53 localization in H1299 cells.** H1299 cells expressing FLAG-tagged FLp53 alone or in combination with V5-tagged isoforms Δ133p53 and Δ160p53 at a ratio of 1:5. To rule out the possibility that the tags might affect protein localization, cells expressing the V5-tagged FLp53 was also analyzed. Cell nuclei were visualized with DAPI (blue). The co-aggregation of isoforms with FLp53 in cytoplasm (white arrows) and nucleus (bright green arrows) are indicated. Scale bar, 50 μm.

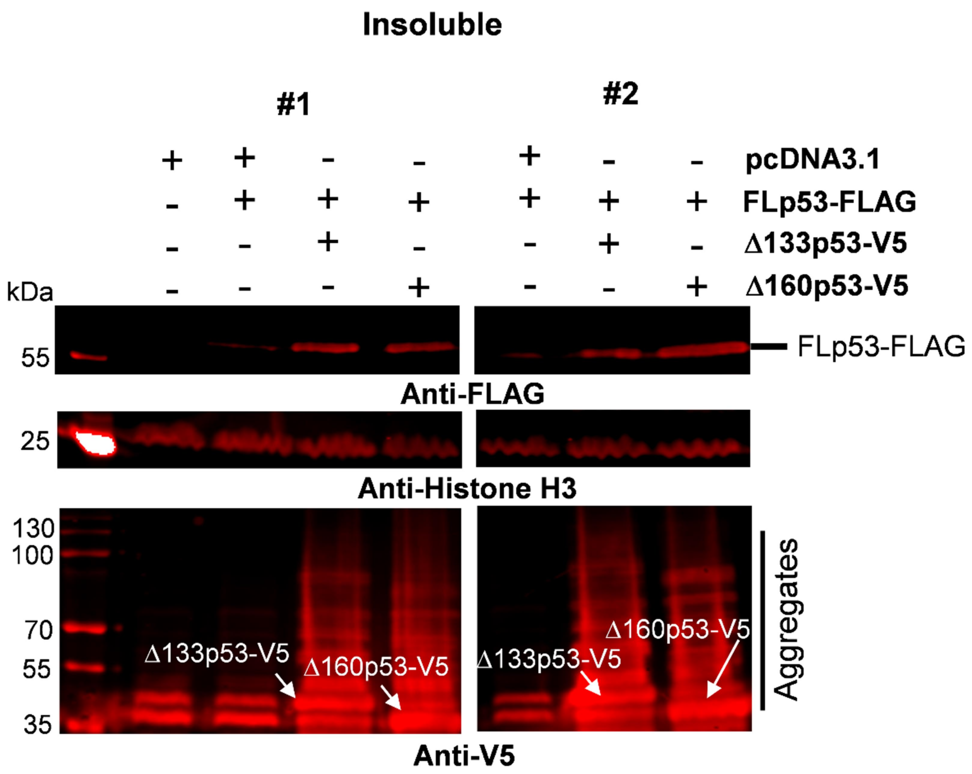


**Supplementary Figure 5. Western-blot analysis of insoluble nuclear fraction of H1299 cells transfected with the FLAG-tagged FLp53, and V5-tagged Δ133p53 or Δ160p53 at a ratio of 1:5.** The insoluble nuclear fraction was run in SDS-PAGE and blotted with Anti-FLAG and Anti-V5 antibodies. Histone H3 were employed as a nuclear marker. While the Anti-FLAG antibody showed a single band of FLp53-FLAG, multiple bands of diverse molecular weight was seen with anti-V5 antibody signifying the aggregated variants of V5-tagged Δ133p53 and Δ160p53 (indicated by white arrows, as well as marked in the margin).

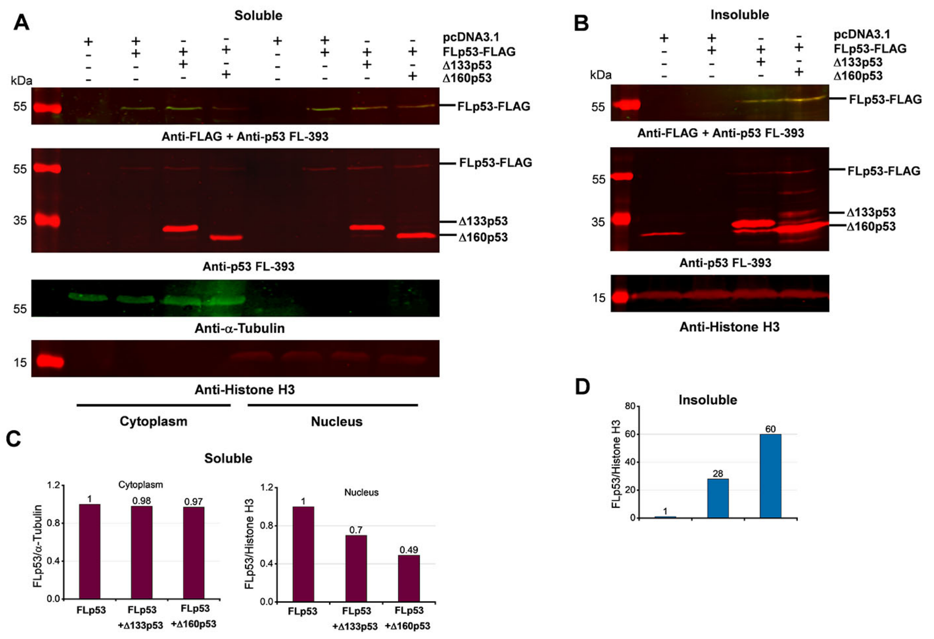


**Supplementary Figure 6. Induction of FLp53 aggregation by Δ133p53 and Δ160p53 isoforms.**

**(A-B)** Western blot analysis of soluble cytoplasmic and nuclear subcellular fractions **(A)** and insoluble nuclear fraction **(B)**. Biochemical fractionation of H1299 cells transfected with the FLAG-tagged FLp53 and untagged Δ133p53 or Δ160p53 at a 1:5 ratio. **(C)** Quantitative bar graph summarizing the levels of the soluble FLp53-FLAG protein in the cytoplasmic and nuclear fractions. The FLp53-FLAG protein levels were normalized to the corresponding fractionation marker (α-Tubulin or histone H3). Relative accumulation of the FLp53-FLAG in Δ133p53 or Δ160p53 expressing cells is shown after considering the FLp53-FLAG protein expressing sample as 1. **(D)** Quantitative bar graph summarizing the levels of the FLp53-FLAG protein in the insoluble nuclear fraction relative to the histone H3. Data normalization as in panel C.

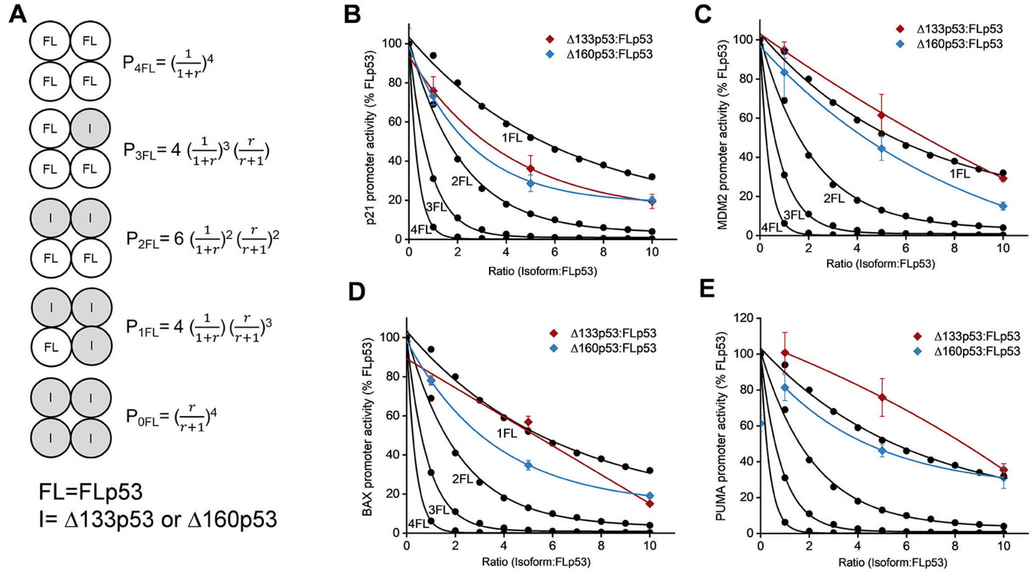


**Supplementary Figure 7. Predictive inhibitory capacity of Δ133p53 and Δ160p53 isoforms in hetero- or homo-tetrameric complexes.**

**(A)** Mathematical modeling of tetramer formation probabilities (adopted from Chan *et al.*[7]). The probabilities (P) of tetramer formation involving different combinations of FLp53(FL) and its isoforms (I) Δ133p53 or Δ160p53 are expressed. The ratio r is defined as the concentration of Δ133p53 or Δ160p53 relative to the concentration of FLp53. **(B-E)** Prediction of isoform/FLp53 hetero-tetramer formation based on the data from Figure 4. The promoter activity of p53 target genes p21 (B), MDM2 (C), BAX (D), and PUMA (E) was shown. For PUMA**,** promoter activities are normalized to the activity when Δ133p53 and FLp53 are co-transfected at a 1:1 ratio. The predicted inhibition curves (black dots) are derived from the tetramer formation probabilities shown in panel A. We hypothesize that the transcription factor is fully active only when it forms a tetramer consisting of 4FL, then the final activity A equals $P_{4FL}$. Thus, $A_{4FL} = P_{4FL}$; $A_{3FL} = P_{4FL} + P_{3FL}$; $A_{2FL} = P_{4FL} + P_{3FL} + P_{2FL}$; $A_{1FL} = P_{4FL} + P_{3FL} + P_{2FL} + P_{1FL}$. The experimental inhibition curves with Δ133p53 (red) or

Δ160p53 (blue) were all above the theoretical inhibition curves for two isoform molecules per tetramer. Our analyses indicate that the Δ133p53 or Δ160p53 isoforms can exert inhibitory effect on FLp53 function only when present in the tetramer in higher proportion than FLp53.
